# *N*^4^-Hydroxycytidine/Molnupiravir Inhibits RNA-Virus Induced Encephalitis by Producing Mutated Viruses with Reduced Fitness

**DOI:** 10.1101/2023.08.22.554316

**Authors:** Durbadal Ojha, Collin S. Hill, Shuntai Zhou, Alyssa B. Evans, Jacqueline M. Leung, Christine Schneider Lewis, Franck Amblard, Raymond F. Schinazi, Ralph S. Baric, Karin E. Peterson, Ronald Swanstrom

**Affiliations:** Laboratory of Neurological Infections and Immunity, Rocky Mountain Laboratories, National Institute of Allergy and Infectious Disease, National Institutes of Health, Hamilton, MT, USA; Lineberger Comprehensive Cancer Center, University of North Carolina at Chapel Hill, Chapel Hill, NC, USA; Department of Microbiology and Immunology, University of North Carolina at Chapel Hill, Chapel Hill, NC, USA; Center for ViroScience and Cure, Laboratory of Biochemical Pharmacology, Department of Pediatrics, Emory University School of Medicine and Children’s Healthcare of Atlanta, Atlanta, Georgia, USA; Department of Epidemiology, University of North Carolina at Chapel Hill, Chapel Hill, North Carolina, USA; Department of Biochemistry and Biophysics, University of North Carolina at Chapel Hill, Chapel Hill, North Carolina, USA; Department of Microbiology and Cell Biology; Montana State University; Montana, USA

## Abstract

A diverse group of RNA viruses including Rabies, Polio, La Crosse, West Nile, Zika, Nipah, Eastern and Western equine encephalitis, Venezuelan equine encephalitis, Japanese encephalitis, and tick-borne encephalitis viruses have the ability to gain access to and replicate in the central nervous system (CNS), causing severe neurological disease. Current treatment for these patients is generally limited to supportive care. To address the need for a generalizable antiviral, we utilized a strategy of mutagenesis to limit virus replication. We evaluated ribavirin (RBV), favipiravir (FAV) and *N*^4^-hydroxycytidine (NHC) against La Crosse virus (LACV) which is the primary cause of pediatric arboviral encephalitis cases in North America. NHC was more potent than RBV or FAV in neuronal cells. Oral administration of molnupiravir (MOV), the 5’-isobutyryl prodrug of NHC, decreased neurological disease development by 32% following intraperitoneal (IP) infection of LACV. MOV also reduced disease by 23% when virus was administered intranasally (IN). NHC and MOV produced less fit viruses by incorporating predominantly G-to-A or C-to-U mutations. Furthermore, NHC also inhibited two other orthobunyaviruses, Jamestown Canyon virus and Cache Valley virus. Collectively, these studies indicate that NHC/MOV has therapeutic potential to inhibit virus replication and subsequent neurological disease caused by this neurotropic RNA virus.

## Introduction

There is a diverse set of RNA viruses that do not circulate in the human population but when introduced from zoonotic sources result in high rates of mortality. Some of these viruses are capable of crossing into the central nervous system (CNS) and infecting brain cells where they induce severe neurological disease including encephalitis or death. This group includes rabies virus (rhabdovirus), La Crosse virus (peribunyavirus), Nipah virus (paramyxovirus virus), Eastern and Western equine encephalitis virus (togaviruses), Venezuelan equine encephalitis virus (togavirus), West Nile virus (flavivirus), Japanese encephalitis virus (flavivirus), Zika virus (flavivirus), and tick-borne encephalitis viruses (flavivirus), as well as several others (*1, 2*). Infections in the human population are sporadic and unpredictable. Some of these viruses are arboviruses that spread to humans through an insect vector, while others spread through contact with an infected mammalian host. While these represent diverse viruses in terms of their genome structures and replication strategies, they all share the feature of having a small RNA genome tightly packed with essential genes needed for viral replication and host defense evasion.

Most antiviral agents utilize one of two strategies: targeting a viral protein function (such as the viral protease as has been done for HIV-1 and SARS-CoV-2) (*3, 4*) or causing chain termination during viral genome replication (which has been accomplished for HIV-1, hepatitis B and C viruses, SARS-CoV-2, and the herpesviruses) (*5, 6*). Targeting a specific viral protein function results in an inhibitor that is very specific to that virus. Chain terminating inhibitors can be effective if the viral polymerase will incorporate the analog and the incorporated chain terminating nucleotide is not removed by a repair or error correction function of the virus. To date, neither of these antiviral drug development strategies has produced a broadly acting inhibitor that could be used on the diverse set of RNA viruses that cause sporadic infections in humans.

These neurotropic RNA viruses do share one vulnerability, their RNA genomes. Viral replication is error prone for RNA viruses, but the introduction of random mutations is limited to a rate that is below a level that results in overall harm to the viral population in the form of excess mutations (*7*). The antiviral strategy of lethal mutagenesis posits that a quick accumulation of the deleterious mutations in these viruses can go beyond the capacity of natural selection to remove such mutations, leading to the loss of fitness and thus favoring host recovery. Mutagenic nucleosides have the ability to pair ambiguously during viral replication, forming either A-U or G-C base pairs but then reversing between the two inducing a mutation. Two early compounds (ribavirin and favipiravir) that have been used as therapeutics give rise to similar mutagenic nucleotides in the cell that can induce mutations in vitro (*8–12*). However, the levels of drug attained *in vivo* are far below the effective doses needed *in vitro*, making it unclear if these inhibitors can function as effective mutagens to be therapeutic in the context of a replicating RNA virus (*13*). In contrast, the mutagenic nucleoside NHC is far more potent *in vitro* where it is possible to show a clear relationship between antiviral activity and mutation load in the viral genome, and its antiviral activity has been shown against a diverse group of RNA viruses (*14–18*). Its 5’-isobutyryl prodrug, MOV is active in mouse models for treating respiratory virus infections and is being used to treat SARS-CoV-2 in humans (*19, 20*).

Given that all RNA viruses share the vulnerability of small RNA genomes packed with essential genes, we reasoned that a potent mutagenic nucleoside such as NHC could be active against neurotropic RNA viruses. To test this idea, we selected La Crosse virus (LACV), a bunyavirus, to assess the activity of NHC *in vitro* and the utility of its prodrug MOV in a neuropathogenic mouse model. LACV is the leading cause of pediatric arboviral encephalitis in North America (*2*) and representative of the larger group of neurotropic viruses. We found that both NHC and MOV showed promising therapeutic potential in multiple neuronal cell lines and two different mouse models by inducing less fit and mutated virus.

## Results

### Effect of mutagenic nucleoside analogs on production of infectious LACV in multiple cell lines

We evaluated ribavirin (RBV), favipiravir (FAV) and NHC against LACV infection in Vero cells for inhibition of virus production using six concentrations, ranging between 1 μM to 300 μM for RBV and FAV, and 0.1 μM to 30 μM for NHC. A 10-fold difference in drug concentration was necessary between NHC and RBV or FAV due to higher cytotoxicity (Table S1). As expected, all three drugs induced a significant reduction in viral titers after viral replication at the higher concentrations of the inhibitors (Fig. 1a). The median effective concentration (EC_50_) for NHC was 0.57 μM, which was roughly 100-fold lower than the EC_50_ of either RBV or FAV, which were 42.7 μM and 58.3 μM, respectively (Fig. 1d).

**Figure 1:**
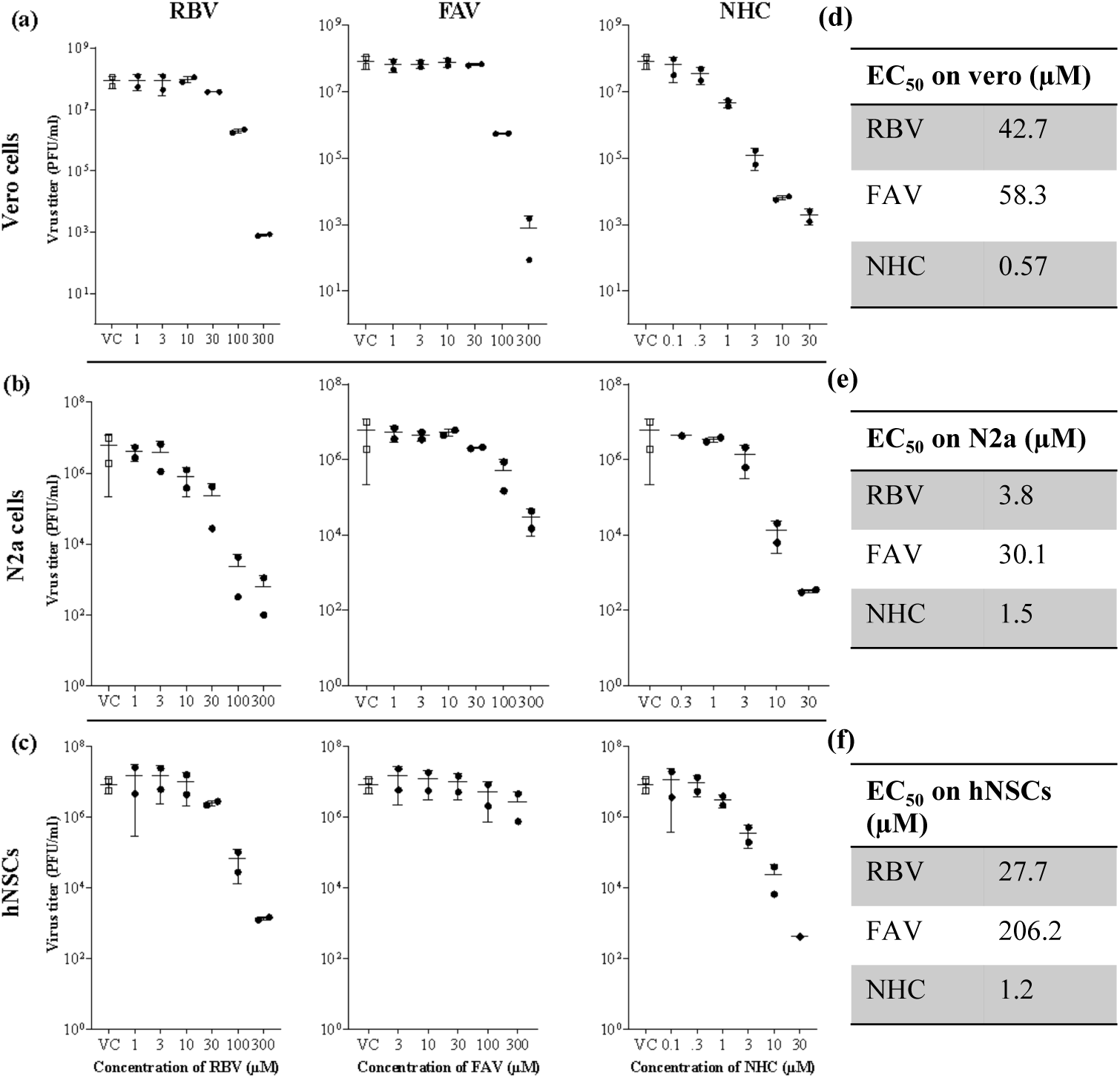
Treatment with NHC inhibited LACV replications more potently than RBV or FAV in multiple cell types. (a-c) Dose-dependent antiviral activity of RBV, FAV and NHC. LACV infected cells were treated with different concentrations of either RBV or FAV or NHC or DMSO (V=vehicle control; VC). At 24 hours post infection (hpi), virus titer was measured from cell supernatant by plaque assay. Each data point represents the number of infectious viruses present in an individual well and are combined from two independent experiments. (d-f) The EC_50_ (median effective concentration) was determined by extrapolating the dose-response curve of either RBV or FAV or NHC in (d) Vero, (e) N2a cells, and (f) hNSCs.

As neurons in the CNS are the primary site of virus replication and tissue damage in humans and in our experimental animal models, we further evaluated each of these nucleoside/nucleobase analogs in rodent-derived neuroblastoma Neuro-2a cells (N2a) and human neural progenitor stem cells (hNSCs). In these cells, NHC and RBV markedly inhibited LACV replication, with an EC_50_ of 1.5 μM in N2a cells and 1.2 μM in hNSC cells for NHC and an EC_50_ of 3.8 μM in N2a cells and 27.7 μM in hNSC cells for RBV (Fig. 1e and 1f). FAV had a moderate effect on LACV replication in N2a cells and a minimal effect in hNSCs, with EC_50_s of 30.1 μM and 206.2 μM, respectively (Fig. 1e, and 1f). This correlated with our recent results showing FAV inhibited LACV replication in Vero cells, but not in neuronal cells (*21*). Overall, NHC was approximately 3 to 170 times more potent than RBV and FAV, which was consistent with our previous studies that NHC is significantly more potent than FAV or RBV against SARS-CoV-2 (*18*). Normal biosynthesis of precursors for RNA synthesis results in nucleoside monophosphates, thus each of these inhibitors must enter a salvage pathway to become biologically active and progress to a nucleoside triphosphate; differences in metabolic activity of these salvage pathways in different cells could account for the varying values of EC_50_ values since the effect of mutations on the virus would be expected to be the same irrespective of cell type.

### Effect of NHC/MOV on LACV-induced neurological disease in a highly pathogenic mouse model

LACV infection of 23-24 day old C57BL/6 mice by intraperitoneal (IP) inoculation of LACV (10^3^ pfu/mice) results in lethal viral encephalitis by 5-7 days post infection (dpi). We tested whether NHC could inhibit LACV induced encephalitis (LACV-E) in mice, by treating mice starting on the day of infection or at 3 dpi, with continued treatment to 10 dpi. The choice of a treatment group starting 3 dpi was based on this timepoint showing the presence of virus in the CNS (*22*). The prodrug of NHC, MOV was given to the mice by oral gavage (300 mg/kg) twice each day. The animals were euthanized with the onset of clinical signs of LACV-induced neurologic disease. Both treatment initiation times with MOVshowed modest efficacy in decreasing the incidence of neurological disease, but this still represents a significant difference from the vehicle-treated mice (Fig. 2a). As can be seen, all of the untreated mice succumbed to viral encephalitis within the first 7 days of infection. When treatment with MOV was started on the day of infection, about 32% of the mice survived the infection (Fig. 2a). When treatment was delayed until 3 dpi, only 15% of the mice survived (Fig. 2a). Thus, MOV was active as an antiviral *in vivo* providing partial protection in this highly pathogenic model.

**Figure 2.**
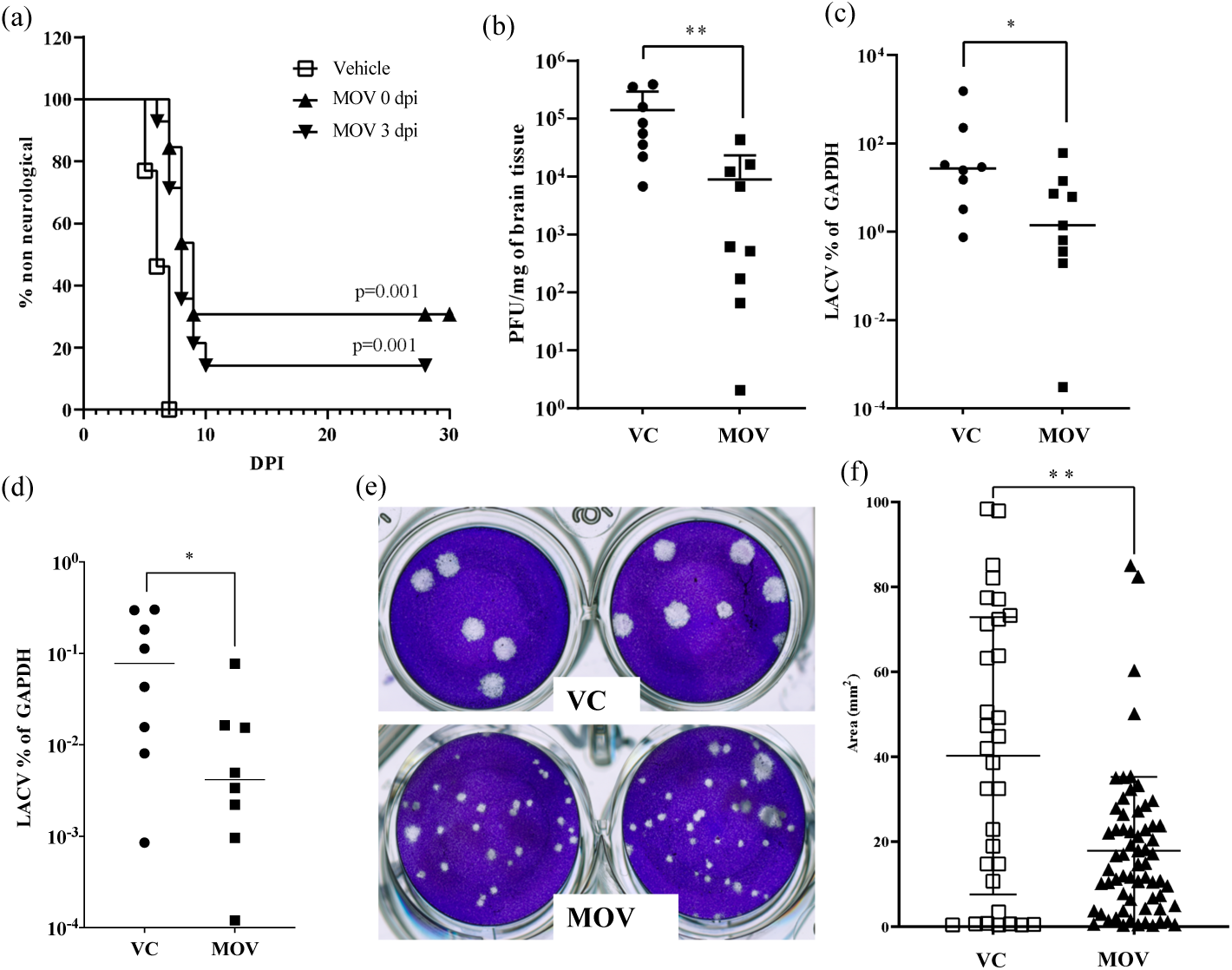
MOV treatment increases survival of LACV-infected mice by inducing less fit and mutated virus in brain tissue. (a) Male and female C57BL/6 mice at the age of day 23 to 25 were infected with LACV (1 x 10^3^ PFU/mice) IP. Mice were treated orally with MOV (300 mg/kg) twice daily until the onset of clinical signs or 10 dpi, starting at day 0 dpi (2 hrs. prior to infection) in one group and 3 dpi in another group. Kaplar-Meier analysis was used to calculate the difference of clinical disease and statistical analysis. (b-f) To quantify the virus titer and plaque size in the CNS and lymph node, tissues were removed at 5 dpi. (b) Infectious virus was quantified using plaque assay and (c, d) viral-RNA by qRT-PCR from (c) brain and (d) lymph node. Each symbol is shown as individual animals from two different independent experiments. (e) Smaller plaque sizes were observed in viral plaque assays of brain tissue from mice receiving MOV treatment compared to mice from the vehicle treated group (VC). (e) To quantify the size of plaques (surface area) formed with and without MOV treatment, wells with clearly isolated plaques were digitized and imported into FIJI for segmentation and analysis. Statistical comparison in (b-d) and (f) were done by using Mann–Whitney U test.

To confirm the antiviral activity of NHC/MOV, we analyzed brain tissue for virus titer using a plaque assay and for viral RNA using qRT-PCR; we also analyzed the presence of viral RNA in lymph nodes using qRT-PCR. Infectious virus titer (Fig. 2b) and the amount of viral RNA (Fig. 2c) in brain tissue and in lymph nodes (Fig. 2d) were significantly decreased in MOV-treated mice. In both the vehicle control and the treated animals, the amount of viral RNA was approximately 100-fold lower in the lymph nodes than in brain tissue, emphasizing the neurotropic nature of this virus. Thus, MOV-mediated reduction of LACV-induced neurological disease was associated with a decreased amount of virus in the periphery and in the CNS.

### MOV treatment induces less fit and mutated LACV in mouse brains

In addition to reduced virus titers in the brain tissue of MOV-treated mice, we noted that the viral plaques from virus isolated from the treated animals appeared to be on average smaller than the plaques from the virus isolated from control animals (Fig. 2e). This could be the case if the mutated genomes were still infectious but carried deleterious mutations. Therefore, we quantified individual plaque size for viruses isolated from vehicle-treated vs MOV-treated animals (Fig. 2f). The average plaque size for virus exposed *in vivo* to NHC was significantly smaller than virus not exposed to NHC. Thus, replication *in vivo* in the presence of NHC resulted not only in reduced virus titer, but also reduced fitness of the residual virus that retained infectivity.

### NHC antiviral activity correlates with mutation density within the LACV genome

NHC functions as a mutagenic ribonucleoside analog with the ability to reversibly base pair as either uridine (U) or cytidine (C) due to tautomerization. To examine this mechanism of action, we sequenced LACV genomes after replication in Vero cells. We used the Primer ID adaptation of a unique molecular identifier (multiplexed to cover several regions of the LACV genome - MPID). The use of Primer ID error correction of each sequenced viral genome provided increased accuracy to detect mutation density above the background error rate of 1 in 10,000 nucleotides sequenced (*23–25*). Using this approach, we were able to establish a correlation between reduced virus titer and mutation density for virus replication in the presence of NHC (Supplemental Fig. 1a). Using the LACV cell culture model, a dose-dependent relationship was observed between the mutation rate of LACV and the concentration of NHC, with similar findings in Vero, hNSCs, and N2a cells (Supplemental Fig. 1b). Additionally, when looking at the mutation profile of the virus cultured with NHC, we saw the greatest increases in C-to-U and G-to-A transition mutation rates, which is consistent with our previous observation on the effect of NHC on the infectivity of multiple coronaviruses (*17, 18*). When we examined the mutation density for LACV grown in Vero cells in the presence of RBV or FAV, we did not see a significant increase in mutation rates except in the highest concentrations of RBV (100 μM) or FAV (100 μM) were used, indicating that NHC is markedly more potent (Supplemental Fig. 1a).

To confirm that the mechanism of action of NHC *in vivo* was also through the mechanism of mutagenesis, we sequenced viral RNA in the isolated tissue of mice that were infected with LACV and treated with MOV. In this model, the total mutation rate of the virus was increased by 2.8 per 10,000 bases in the presence of MOV when compared to vehicle controls (with the mutation rate averaged across the different genomic regions/amplicons sequenced). Specifically, the C-to-U transition rate was increased by 9.1 per 10,000 bases and the G-to-A transition rate was increased by 5.2 per 10,000 bases (Fig. 3a), consistent with our findings with the LACV cell culture model. When we plotted the overall mutation rate of LACV exposed to MOV and LACV copy numbers in the murine brain tissues measured by qRT-PCR, we found a negative logarithmic relationship, which supports the lethal mutagenesis mechanism of MOV/NHC *in vivo* (Fig. 3b). We estimate that each single mutation increase per 10,000 nucleotides resulted in a 4.4-fold decrease in viral load in brain tissue.

**Figure 3.**
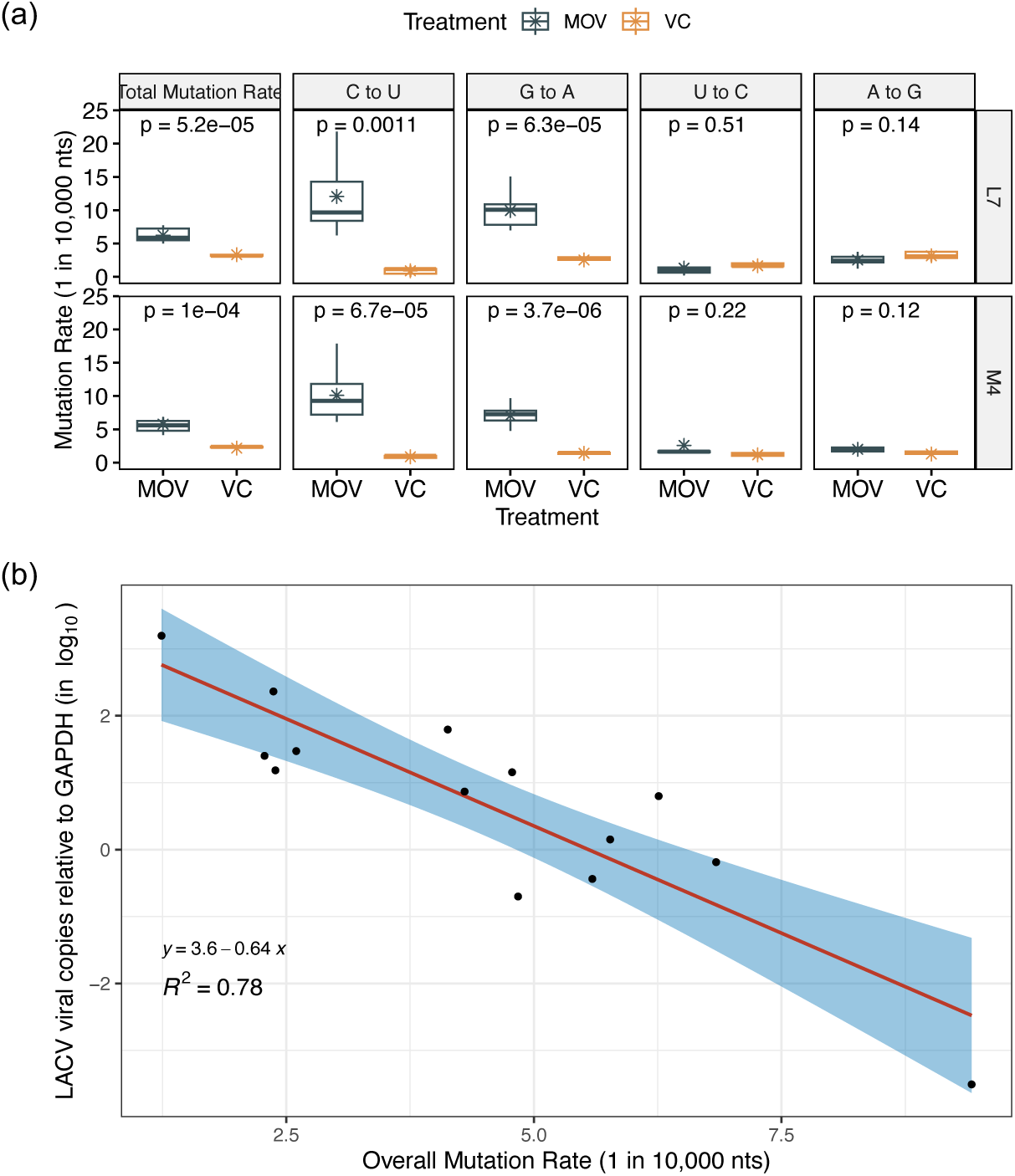
MPID-NGS of the mechanism of action of MOV in LACV mouse model. (a) Mutation rate of LACV in the brain tissues after MOV or vehicle control (VC) treatment. (b) Correlation of the overall mutation rate and LACV copy numbers in the brain tissues measured by qRT-PCR.

### Antiviral effect of MOV given orally during LACV CNS infection

We have previously shown that the LACV induces vascular leakage which promotes viral neuroinvasion (*22*). It is not known if MOV crosses the blood-brain barrier (BBB), although its ability to reduced LACV-induced disease (Fig. 2) suggests either it has ability to cross the BBB (consistent with significant inhibition of disease after starting treatment at 3 dpi) or that treatment produced mutated virus in the periphery that crossed the BBB but with limited disease potential. To address this, we infected the mice with LACV (10^2^ PFU/ml) *via* an intranasal (IN) inoculation, which provides a direct route of access to the CNS through olfactory sensory nerves. Mice were treated orally with MOV or vehicle control starting on the day of infection. LACV-infected mice treated with vehicle control had a clinical disease rate of about 45% (Fig. 4a). MOV treatment decreased the incidence of neurological disease to about 23%, or about a 50% reduction in disease in the less pathogenic model (Fig. 4a). Infectious virus titer was slightly, but not significantly decreased with some variability between mice (Fig. 4b). However, viral RNA expression (Fig. 4c) was significantly decreased in the MOV-treated mice. When we sequenced the LACV viral RNA from the brain samples using MPID-NGS, we found increases in mutation rates similar to that with IP-inoculated and treated mice, although the levels were reduced about two-fold (Fig. 4d). MOV increased both C-to-U mutations and G-to-A mutations compared to vehicle control, with C-to-U and G-to-A mutations increasing by 4.3 per 10,000 bases (Fig. 4d). While these trends are the same as what is seen with virus sequenced after treatment in the IP infection model (Fig. 3), the two-fold difference in mutation density in the treatment group reduced the number of comparisons that reached statistical significance (Fig. 4d).

**Figure 4.**
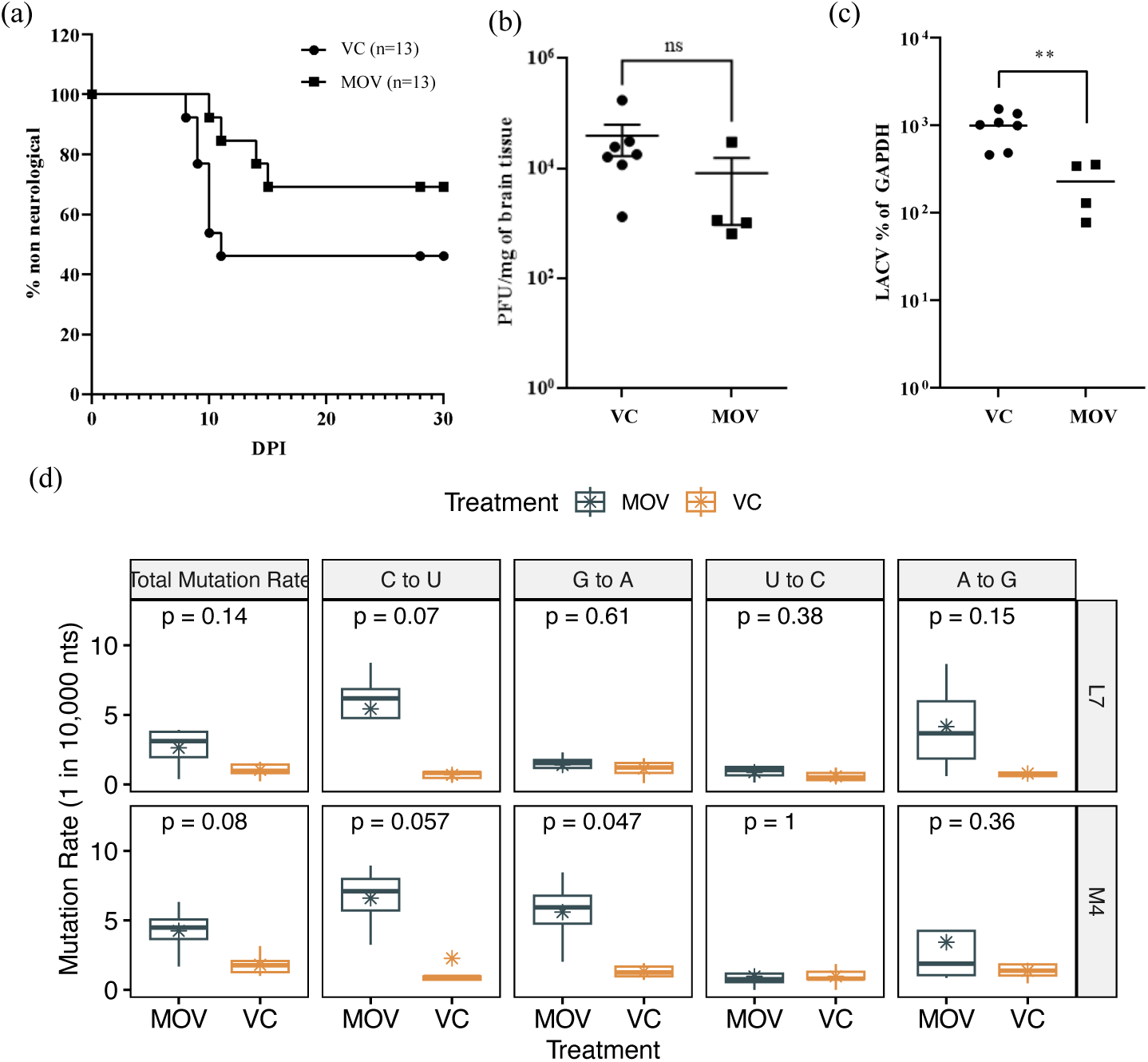
MOV treatment increases survival after infection with through an IN route. (a-c) A mix of both male and female C57BL/6 mice at the age of day 23 to 25 were infected with LACV (1 x 10^2^ PFU/mice) through an IN route. Mice were treated orally with molnupiravir (300 mg/kg; DOSING ROUTE?) twice daily until the onset of clinical signs or 10 dpi, starting at day 0 (2 hrs. prior to infection) and followed for development for clinical disease. Kaplar-Meier analysis were used to calculate the difference of clinical disease and statistical analysis (a). Brain tissue was harvested after animals showed clinical signs of the disease. To quantify the virus titer of these clinical mice we used (b) a plaque assay for quantifying infectious virus titer and (c) qRT-PCR for viral-RNA. (d) Mutation density of LACV in brain tissue in presence or absence (vehicle) of NHC was measured by MPID-NGS in two sequenced regions (L7 and M4). Statistical comparisons in panel (b-d) were performed using Mann-Whitney U test.

### Serial passage of LACV in presence of NHC to select for drug resistance virus

LACV was passaged in Vero cells in the presence of DMSO vehicle carrier (control) or with increasing concentrations of NHC from 0.1 to 0.8 µM (Fig. 5a). After 20 passages (five at each of the increasing NHC concentrations), cell supernatants were harvested. The antiviral activity of NHC was determined in Vero cells for both the control virus and the NHC-selected virus. The EC_50_ values were the same (0.5 μM and 0.49 μM) against both the control and selected viruses (Fig. 5b and 5c). Thus, serial passage of LACV in the presence of NHC using this protocol failed to select for virus with reduced sensitivity to NHC. In addition, the mutagenic effect of NHC can be inferred in that the supernatant virus titer at the end of the selection period was approximately 15-fold lower than for the control passaged virus, and with reduced plaque size (data not shown).

**Figure 5:**
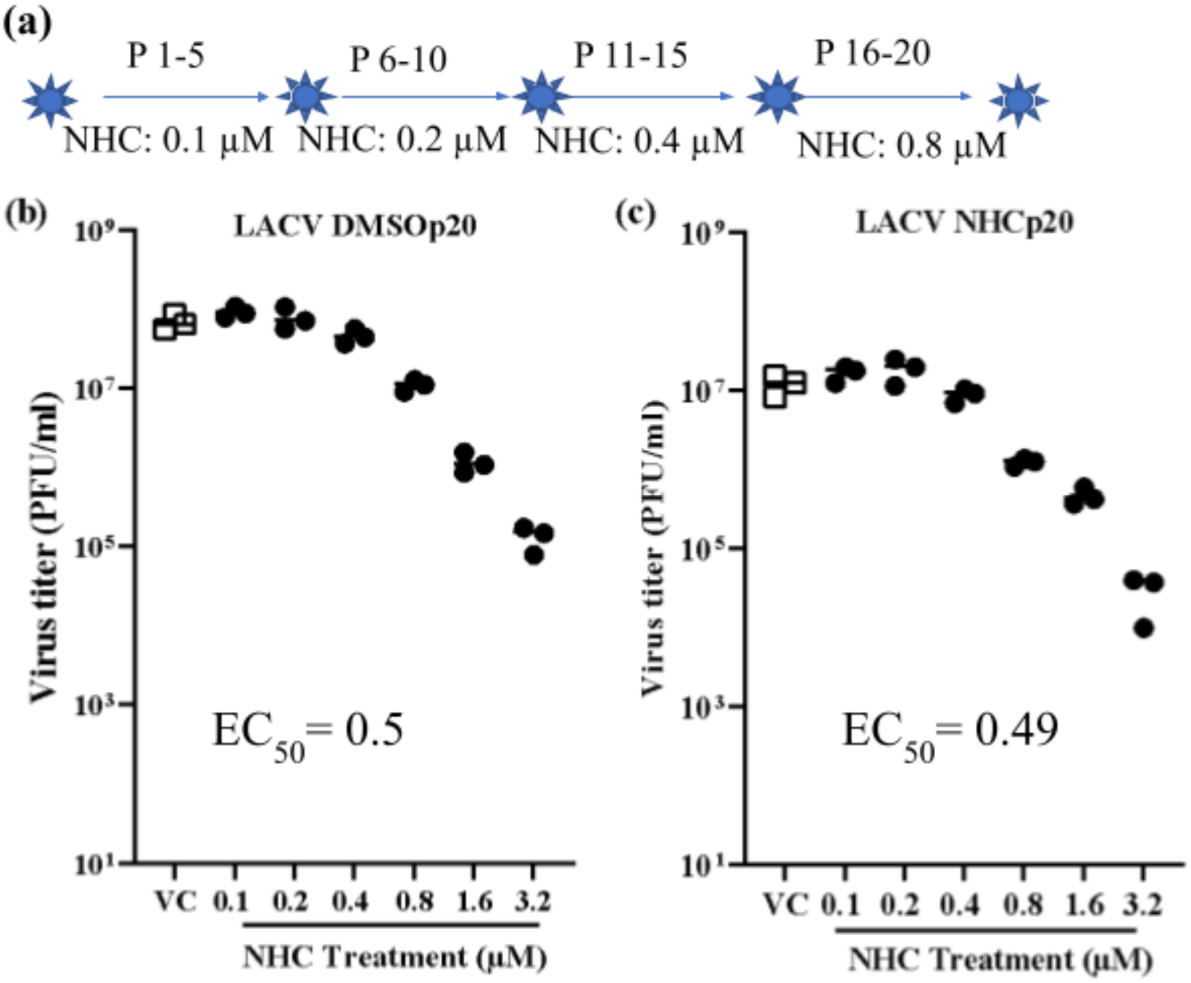
Serial passage of LACV in the presence of increasing NHC concentrations to select for resistance. (a) Study protocol of serial passaging of LACV in presence of DMSO or increasing concentrations of NHC (0.1 to 0.8 µM), with virus being passaged five times at each concentration. (b-c) Dose-dependent antiviral activity of NHC against serially passaged LACV. Passaged viruses were used to infect Vero cells treated with different concentrations of NHC or with DMSO (VC). At 24 hr post infection (hpi), virus titer was measured from cell supernatant using a plaque assay. Each data point represents the number of infectious viruses present in an individual well from independent experiments.

### Effect of NHC on other California Serogroup (CSG) orthobunyaviruses

We also evaluated the ability of NHC to inhibit the replication of two other orthobunyaviruses, Cache Valley virus (CVV) and Jamestown Canyon virus (JCV). JCV belongs to the same serogroup as LACV, but causes neurological disease in both children and adults. CVV is a member of the Bunyamwera serogroup and is one of the most common orthobunyavirus in the United States. Infections were done following the same protocol as for Fig. 1. NHC treatment efficiently reduced CVV and JCV infectious virus production in Vero cells with EC_50_ values of 0.25 and 0.3 μM, respectively (Fig. 6). Thus, NHC inhibited virus replication for multiple orthobunyaviruses in different serogroups.

**Figure 6:**
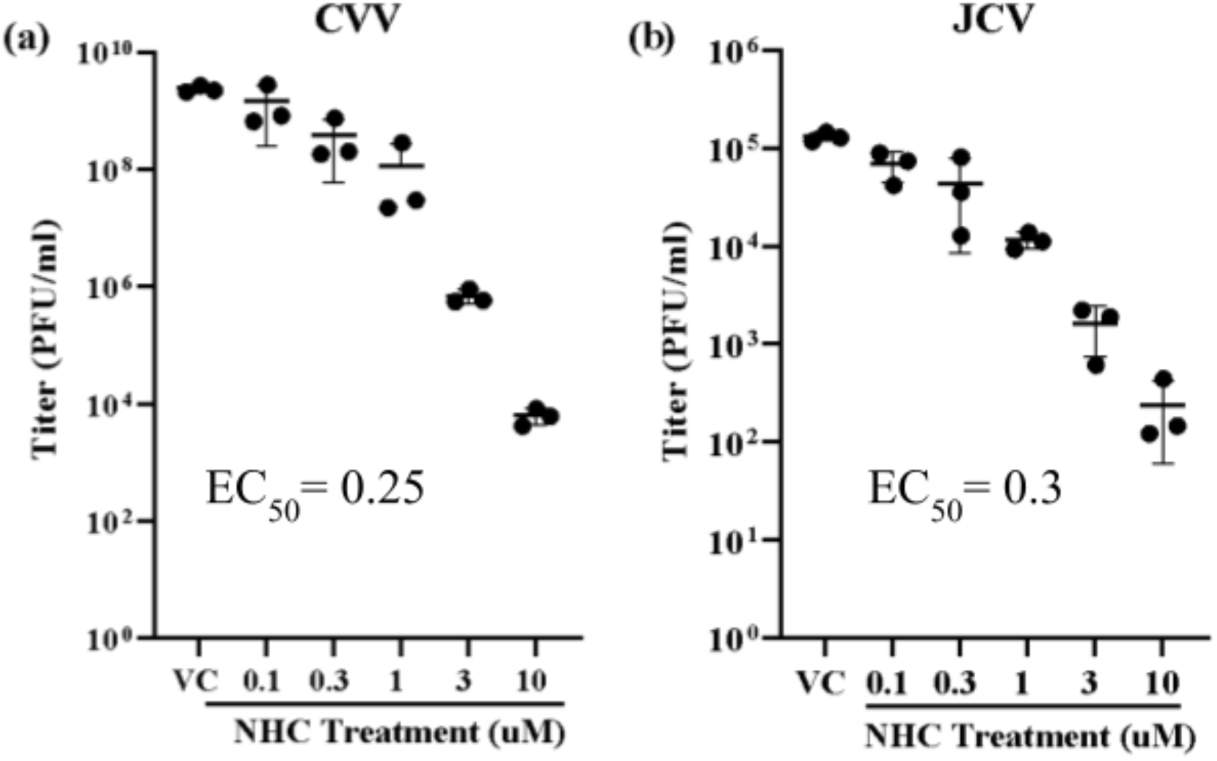
Dose-dependent activity of NHC on (a) CVV and (b) JCV replication. Virus infected cells were treated with different concentrations of NHC or DMSO (VC). At 24 hpi, virus titer was measured from cell supernatant by plaque assay. Each data point represents the PFU/ml from an individual experiment, and three independent experiments were performed.

## Discussion

In this study, we used LACV, a member of the California Serogroup (CSG) of orthobunyaviruses, as a representative neurotropic RNA virus to study the antiviral effect of mutagenic nucleoside NHC/MOV in a viral encephalitis model. After comparative analysis of three mutagenic nucleoside analogs against LACV replication in different neuronal cell lines, we found that NHC was more effective than RBV and FAV at inhibiting virus replication. We also showed *in vivo* efficacy, in that treatment with the NHC prodrug MOV increased survival of LACV-infected mice and induced less fit and mutated virus in brain tissue. Although we did not measure uptake or intracellular states of these nucleotide precursors, it seems likely that the differences in potency between the three mutagenic nucleoside/nucleobase analogs represent differences in cellular uptake, initial metabolism to the nucleoside triphosphate, and/or incorporation by the viral polymerase. The availability of a suitable animal models is essential to evaluate potential drug target. In this study, we used weanling C57Bl/6 mouse model which mimics clinical LACV disease pathology in humans. Weanling C57Bl/6 mice are susceptible to LACV infection either through the IP (peripheral) or IN (direct access to the brain) route of inoculation. We initially used the IP route of inoculation which mimic arboviruses virus transmission to humans by the bite of an infected insect vector. MOV treatment significantly increased survival of LACV-infected mice and generated mutated and less fit virus in the brain (Fig. 2). In a second experiment, we utilized the IN route of inoculation to determine if the drug when given orally was able to display an antiviral effect for virus replication in the brain. In this second experiment, there was a trend of increased survival with MOV treatment which was consistent with our previous IP inoculated model. Even in this stringent viral encephalitis model, we were still able to detect increased mutagenesis of the viral genome in the brain (Fig. 4d) suggesting the presence of the drug within the CNS even after oral administration. It is possible that intrathecal administration could provide a more efficacious dosing strategy in treating cases of viral encephalitis.

NHC has been demonstrated to be an active antiviral against a number of RNA viruses including in VEEV, SARS-CoV-2, MERS-CoV, Ebola virus, LACV (and other California serogroup orthobunyaviruses), and others (*14–18*). It is proposed that the mutagenic effect of MOV/NHC is due to the presence of the 4-oxime group, which can undergo tautomerization between an amino and an imino form to allow MOV/NHC to ambiguously base pair with either adenosine or guanosine depending on its tautomeric form (*26, 27*). Since RNA viruses must undergo both plus strand and minus strand RNA synthesis where NHC can be incorporated, both G-to-A or C-to-U mutations appear in the plus/coding strand. These findings are consistent with the mechanism of action for MOV/NHC in LACV being through lethal mutagenesis, whereby increasing the mutation rate of the virus past its error threshold results in decreases in the viral population mean fitness and eventual viral population collapse due to the accumulation of deleterious mutations (*28*). This is further supported by our findings in Fig. 3b, where we found a negative logarithmic relationship between the overall mutation rate of LACV exposed to MOV and LACV copy numbers in the murine brain tissues measured by qRT-PCR. While lethal mutagenesis is the end point of excessive mutations, in practice the treatment likely is creating less fit virus and reduced virus titers that allow host immune mechanisms to contain the infection. The potency of NHC is likely due to its close similarity to cytidine, allowing it to be readily metabolized and incorporated. This may also be the reason we were unable to select for resistance to NHC (Fig. 5), although selection for low-level resistance to NHC has been reported for murine hepatitis virus (MHV) (*29*).

Ribavirin has been previously studied against LACV, including in phase I, IIa, and IIb clinical trials in pediatric patients with severe LACV-E, however its efficacy remains unclear. While ribavirin has previously shown strong antiviral activity against LACV in cell culture, its efficacy in clinical trials was poor due to reduced penetration into the CNS, resulting in subtherapeutic CSF concentrations when ribavirin was administered *via* IV (*13, 30*). Additionally, when higher doses of ribavirin were given to increase CSF concentrations, not only did the levels of ribavirin still fail to consistently reach therapeutic levels, but the higher doses were associated with higher rates of adverse events, including hyperammonemia and pancreatitis, thus suggesting a narrow therapeutic window for ribavirin in the treatment of LACV-E (*13*). Off-target effects on the host have been suggested to contribute to toxicity associated with RBV treatment and may also contribute to the modest observed antiviral effects (*13*); the weak potency of RBV (and FAV) as viral mutagens compared to NHC likely will make NHC a more potent antiviral for this class of drugs with broad activity against RNA viruses. MOV has also been shown to be well tolerated in clinical trials for treatment of adults with COVID-19 with minimal adverse events grade 3 or above and thus potentially providing a broader therapeutic window when compared to ribavirin (*13, 31*).

Our study is the first *in vivo* therapeutic model of encephalitic RNA virus infection using mutagenic compounds as direct-acting antiviral agents. The results highlight the potential of using such compounds, particularly NHC/MOV in the treatment of other CNS infection caused by RNA viruses. Additionally, further study is needed to examine intrathecal administration of MOV (or NHC) for treatment of encephalitic RNA virus infection. While our mouse model suggests that MOV is able to cross the BBB when given systemically, intrathecal administration may also enhance the levels of drug within the CNS by direct CSF introduction. Furthermore, the intrathecal delivery of MOV offers the potential to mitigate systemic drug toxicity-related adverse effects. This approach could potentially minimize the exposure of high-turnover cells at greatest risk of genotoxicity throughout the body, such as GI tract epithelial cells or hematopoietic stem cells, reducing the risk for the development of germline mutations in men and of cancer.

## Materials and Methods

### Compounds

For all cell culture experiments, *N*^4^-β-hydroxycytidine (NHC) was obtained through the NIH AIDS Reagent Program, Division of AIDS, NIAID, NIH, as a solid to be mixed in DMSO (originally provided by Dr. Schinazi. For all mouse model experiments, MOV was supplied by the Schinazi laboratory at Emory University, Atlanta, Georgia, USA.

### Cells

Vero cells (African green monkey kidney cells, ATCC) were used for the initial screening study and also for viral titer determination by plaque assays, virus maintenance and virus stock production. N2a cells (ATCC CCL-131) were grown and maintained in Dulbecco’s modified Eagle medium (DMEM) containing 4,500 mg/L D-glucose, 4 mM L-glutamine and 1mM sodium pyruvate supplemented with 10% fetal bovine serum (FBS). Human (H9) embryonic stem cells-derived hNSCs (Thermo Fisher Scientific) were cultured in KnockOut™ D-MEM/F-12 media (Thermo Fisher Scientific).

### Antiviral efficacy study by quantified virus titration and EC50 determination

Cells were seeded in 24-well plates (1 × 10^5^ cells/well) and infected with LACV (0.1 MOI). At 1 hr post infection (hpi), medium was replaced with fresh medium containing 6 different concentrations for each of the drugs. RBV and FAV were tested at concentrations of 1, 3, 10, 30, 100, and 300 μM, while NHC was tested at concentrations of 0.1, 0.3, 1, 3, 10 and 30 μM. DMSO was used as vehicle control. After 24 hpi, the cell supernatant was harvested, centrifuged to eliminate cell debris and stored at −80 °C for sequencing (described below) and plaque assay following the method of Ojha et al. 2021 (*21*). Quantified virus titers were plotted to represent antiviral efficacy. The EC_50_ (Median effective concentration) was determined by extrapolating dose-response curve using GraphPad Prism 8 and 9.

### Quantification of LACV viral RNA genome copy number by qRT-PCR

Viral RNA was isolated using the Zymo Direct-zol RNA MiniPrep Kit (Zymo Research) per the manufacturer’s directions. First-strand complementary DNA (cDNA) was generated using iScript reverse transcriptase (Bio-Rad). qRT-PCR was performed using PowerUp SYBR Green Master Mix (ThermoFischer) on QuantStudio 6 (Applied Biosystems). Primers used for the reaction were synthesized by Integrated DNA Technologies (IDT) to quantify LACV viral RNA, targeting the S segment of the genome: forward primer 5’-ATTCTACCCGCTGACCATTG-3’, reverse primer 5’-GTGAGAGTGCCATAGCGTTG-3’. Primers for mouse Gapdh are forward primer 5’-AACGACCCCTTCATTGAC-3’ and reverse primer 5’-TCCACGACATACTCAGCA-3’.

### Primer ID Sequencing of viral RNA

We used a Primer ID (PID) library preparation approach with Illumina MiSeq next-generation sequencing (NGS) to examine the mutagenic effects of these drugs on viral RNA allowing deep coverage of the viral population while maintaining a low method error rate (1 in 10,000 nucleotides) (*23–25*). We designed cDNA primers targeting 372 bp region of the RdRp gene within the L segment (referred as L7 in the study), and included a second amplicon covering a 455 bp portion of the M segment (referred as M4 in the study). Each cDNA primer included a block of random nucleotides (11-bp long) as the primer ID/unique molecular identifier (UMI). Extracted viral RNA, was reversed transcribed using SuperScript III (Thermal Fisher) to make cDNA in a single reaction with the two different cDNA primers. After bead purification, the cDNA was amplified by PCR using a mixture of forward primers and a universal reverse primer, followed by a second round of PCR to incorporate Illumina sequencing adapters and barcodes within the amplicons. After gel-extraction of the PCR product (Qiagen MinElute Gel Extraction kit), the extracted libraries were quantified using a Qubit fluorometer and subjected to Agilent TapeStation analysis. For each sequencing submission, 24 quantified libraries were pooled for MiSeq 300 bp paired-end sequencing at UNC High Through-put Sequencing Facility. Primers for library prep can be found in Table S2. The initial Primer ID data was processed using the Illumina bcl2fastq pipeline (version 2.20.0), and then template consensus sequences (TCSs) were constructed using the tcs pipeline version 2.5.1 (https://primer-id.org/) (*25*). Each type of substitution rate was calculated against the wildtype sequence at the sequenced regions. National Center for Biotechnology Information (NCBI) sequence read archive (SRA) accession numbers are pending.

### LACV encephalitis (LACV-E) mouse model

All animal experiments were performed following the protocols 2019-023-E and 2022-019-E approved by the NIH/NIAID/RML Institutional Animal Care and Use Committee. For this study, a mix of male and female C57BL/6 mice at 23 to 25 days old were used. Mice were divided evenly and infected with LACV at either 1 × 10^3^ PFU by IP route or 1 x 10^2^ PFU by IN route. IP-infected mice were started on treatment with MOV (ROUTE? IP?) at either the date of infection or at 3 dpi; the IN-infected mice were started on treatment on the day of infection. Molnupiravir was diluted in 10% polyethylene glycol (PEG), 2.5% Cremophor RH40 and 87.5% sterile water which was used as vehicle control. For oral administration, 300 mg/kg of MOV twice a day was used for up to 10 consecutive days using a straight 24-gauge ball-tip disposable feeding needle (Instech). Mice were examined twice daily for 28-30 days for clinical signs of LACV-induced neurologic disease, which mostly involved limb paralysis, ataxia, and weakness or repeated seizures. Mice showing one of these clinical symptoms were euthanized for tissue removal and scored as clinical.

### Tissue harvesting for plaque assay

Brain tissue and lymph nodes were harvested from mice with clinical symptoms (as describe above) or at 5 dpi and stored at -80 °C. Half of the sagittal section of the brain tissue was used for plaque assays, and the other half for RNA isolation for qRT-PCR and sequencing. For the plaque assay, brain tissues were homogenized in presence of DMEM using the Bead Mill 24 (Fisher Scientific) homogenizer at 5300 rpm for 25 s. Homogenized samples were centrifuged at 5000 x*g* for 10 min. Supernatants were then diluted in DMEM containing 10% FBS to 10^-1^ to 10^-7^ dilutions and finally subjected to plaque assay (*32*).

### Measurement of plaque size

A digital scanner was used to scan the plaque assay plates. Digitized images were imported into FIJI/ImageJ v1.52n (*33*), and pixel calibration values were calculated based on the well diameter and corresponding digitized image. To quantify plaque size (surface area), wells with clearly isolated plaques were selected for segmentation and analysis. The image lookup table for each well was inverted followed by binarization prior to analysis, with a minimum area of 0.39 mm^2^ to eliminate false positives due to slight variations in crystal violet staining. The measurements were imported into Microsoft Excel for data compilation, and graphed and analyzed using GraphPad Prism v9.3.1. Analysis of plaque size was done only when plaque assays for different virus samples were done in parallel.

### Statistical Analysis

All post-study statistical analysis were conducted using GraphPad Prism v.8.02 and v.9.3.1. All P values are represented as *P <0.05, **P <0.01, ***P <0.001, and ****P<0.0001. Specific analysis information is described in figures legends.

## Acknowledgements

We acknowledge the efforts of the University of North Carolina (UNC) High Throughput Sequencing Facility.

## Literature Cited

1. J. T. Gaensbauer, N. P. Lindsey, K. Messacar, J. E. Staples, M. Fischer, Neuroinvasive arboviral disease in the United States: 2003 to 2012. Pediatrics 134, e642–650 (2014).

2. R. A. Soto, M. L. Hughes, J. E. Staples, N. P. Lindsey, West Nile virus and other domestic nationally notifiable arboviral diseases - United States, 2020. MMWR Morb Mortal Wkly Rep 71, 628–632 (2022).

3. C. Flexner, HIV-protease inhibitors. N Engl J Med 338, 1281–1292 (1998).

4. S. Iketani, F. Forouhar, H. Liu, S. J. Hong, F.-Y. Lin, M. S. Nair, A. Zask, Y. Huang, L. Xing, B. R. Stockwell, Lead compounds for the development of SARS-CoV-2 3CL protease inhibitors. Nature Communications 12, 2016 (2021).

5. T. Cihlar, A. S. Ray, Nucleoside and nucleotide HIV reverse transcriptase inhibitors: 25 years after zidovudine. Antiviral Research 85, 39–58 (2010).

6. C. D. Spinner, R. L. Gottlieb, G. J. Criner, J. R. A. López, A. M. Cattelan, A. S. Viladomiu, O. Ogbuagu, P. Malhotra, K. M. Mullane, A. Castagna, Effect of remdesivir vs standard care on clinical status at 11 days in patients with moderate COVID-19: a randomized clinical trial. JAMA 324, 1048–1057 (2020).

7. J. J. Bull, R. Sanjuan, C. O. Wilke, Theory of lethal mutagenesis for viruses. J Virol 81, 2930–2939 (2007).

8. A. B. Janowski, H. Dudley, D. Wang, Antiviral activity of ribavirin and favipiravir against human astroviruses. J Clin Virol 123, 104247 (2020).

9. L. Oestereich, T. Rieger, M. Neumann, C. Bernreuther, M. Lehmann, S. Krasemann, S. Wurr, P. Emmerich, X. de Lamballerie, S. Ölschläger, S. Günther, Evaluation of antiviral efficacy of ribavirin, arbidol, and T-705 (favipiravir) in a mouse model for Crimean-Congo hemorrhagic fever. PLoS Negl Trop Dis 8, e2804–e2804 (2014).

10. J. T. Rankin, Jr., S. B. Eppes, J. B. Antczak, W. K. Joklik, Studies on the mechanism of the antiviral activity of ribavirin against reovirus. Virology 168, 147–158 (1989).

11. E. Vanderlinden, B. Vrancken, J. Van Houdt, V. K. Rajwanshi, S. Gillemot, G. Andrei, P. Lemey, L. Naesens, Distinct effects of T-705 (Favipiravir) and Ribavirin on influenza virus replication and viral RNA synthesis. Antimicrob Agents Chemother 60, 6679–6691 (2016).

12. T. Baranovich, S.-S. Wong, J. Armstrong, H. Marjuki, R. J. Webby, R. G. Webster, E. A. Govorkova, T-705 (favipiravir) induces lethal mutagenesis in influenza A H1N1 viruses in vitro. J Virol 87, 3741–3751 (2013).

13. J. E. McJunkin, M. C. Nahata, E. C. De Los Reyes, W. G. Hunt, M. Caceres, R. R. Khan, M. G. Chebib, S. Taravath, L. L. Minnich, R. Carr, C. A. Welch, R. J. Whitley, Safety and pharmacokinetics of ribavirin for the treatment of la crosse encephalitis. Pediatr Infect Dis J 30, 860–865 (2011).

14. M. Ehteshami, S. Tao, K. Zandi, H. M. Hsiao, Y. Jiang, E. Hammond, F. Amblard, O. O. Russell, A. Merits, R. F. Schinazi, Characterization of β-D-N(4)-hydroxycytidine as a novel inhibitor of Chikungunya virus. Antimicrob Agents Chemother 61, PAGES? (2017).

15. K. Zandi, F. Amblard, S. Amichai, L. Bassit, S. Tao, Y. Jiang, L. Zhou, O. Ollinger Russell, S. Mengshetti, R. F. Schinazi, Nucleoside analogs with antiviral activity against Yellow Fever virus. Antimicrob Agents Chemother 63, e00889–00819 (2019).

16. N. Urakova, V. Kuznetsova, D. K. Crossman, A. Sokratian, D. B. Guthrie, A. A. Kolykhalov, M. A. Lockwood, M. G. Natchus, M. R. Crowley, G. R. Painter, E. I. Frolova, I. Frolov, beta-D-N (4)-Hydroxycytidine Is a Potent anti-alphavirus compound that induces a high level of mutations in the viral genome. J Virol 92, e01965–01917 (2018).

17. T. P. Sheahan, A. C. Sims, S. Zhou, R. L. Graham, A. J. Pruijssers, M. L. Agostini, S. R. Leist, A. Schäfer, K. H. Dinnon, 3rd, L. J. Stevens, J. D. Chappell, X. Lu, T. M. Hughes, A. S. George, C. S. Hill, S. A. Montgomery, A. J. Brown, G. R. Bluemling, M. G. Natchus, M. Saindane, A. A. Kolykhalov, G. Painter, J. Harcourt, A. Tamin, N. J. Thornburg, R. Swanstrom, M. R. Denison, R. S. Baric, An orally bioavailable broad-spectrum antiviral inhibits SARS-CoV-2 in human airway epithelial cell cultures and multiple coronaviruses in mice. Sci Transl Med 12, (2020).

18. S. Zhou, C. S. Hill, S. Sarkar, L. V. Tse, B. M. D. Woodburn, R. F. Schinazi, T. P. Sheahan, R. S. Baric, M. T. Heise, R. Swanstrom, β-D-N^4^-hydroxycytidine inhibits SARS-CoV-2 through lethal mutagenesis but is also mutagenic to mammalian cells. J Infect Dis 224, 415–419 (2021).

19. W. A. Fischer, J. J. Eron Jr, W. Holman, M. S. Cohen, L. Fang, L. J. Szewczyk, T. P. Sheahan, R. Baric, K. R. Mollan, C. R. Wolfe, A phase 2a clinical trial of molnupiravir in patients with COVID-19 shows accelerated SARS-CoV-2 RNA clearance and elimination of infectious virus. Sci Transl Med 14, eabl7430 (2021).

20. A. Jayk Bernal, M. M. Gomes da Silva, D. B. Musungaie, E. Kovalchuk, A. Gonzalez, V. Delos Reyes, A. Martín-Quirós, Y. Caraco, A. Williams-Diaz, M. L. Brown, Molnupiravir for oral treatment of Covid-19 in nonhospitalized patients. New England Journal of Medicine 386, 509–520 (2022).

21. D. Ojha, C. W. Winkler, J. M. Leung, T. A. Woods, C. Z. Chen, V. Nair, K. Taylor, C. D. Yeh, G. J. Tawa, C. L. Larson, W. Zheng, C. L. Haigh, K. E. Peterson, Rottlerin inhibits La Crosse virus-induced encephalitis in mice and blocks release of replicating virus from the Golgi body in neurons. Nature Microbiology 6, 1398–1409 (2021).

22. C. W. Winkler, B. Race, K. Phillips, K. E. Peterson, Capillaries in the olfactory bulb but not the cortex are highly susceptible to virus-induced vascular leak and promote viral neuroinvasion. Acta Neuropathol 130, 233–245 (2015).

23. S. Zhou, C. Jones, P. Mieczkowski, R. Swanstrom, Primer ID validates template sampling depth and greatly reduces the error rate of next-generation sequencing of HIV-1 genomic RNA populations. J Virol 89, 8540–8555 (2015).

24. S. Zhou, C. S. Hill, M. U. Clark, T. P. Sheahan, R. Baric, R. Swanstrom, Primer ID next-generation sequencing for the analysis of a broad spectrum antiviral induced transition mutations and errors rates in a coronavirus genome. Bio Protoc 11, e3938 (2021).

25. S. Zhou, C. S. Hill, E. Spielvogel, M. U. Clark, M. G. Hudgens, R. Swanstrom, Unique molecular identifiers and multiplexing amplicons maximize the utility of deep sequencing to critically assess population diversity in RNA viruses. ACS Infect Dis 8, 2505–2514 (2022).

26. F. Amblard, J. C. LeCher, R. De, S. L. Goh, C. Li, M. Kasthuri, N. Biteau, L. Zhou, Z. Tber, J. Downs-Bowen, K. Zandi, R. F. Schinazi, Synthesis of novel N(4)-hydrocytidine analogs as potential anti-SARS-CoV-2 agents. Pharmaceuticals (Basel) 15, PAGES?? (2022).

27. F. Kabinger, C. Stiller, J. Schmitzová, C. Dienemann, G. Kokic, H. S. Hillen, C. Höbartner, P. Cramer, Mechanism of molnupiravir-induced SARS-CoV-2 mutagenesis. Nat Struct Mol Biol 28, 740–746 (2021).

28. M. Eigen, Error catastrophe and antiviral strategy. Proc Natl Acad Sci U S A 99, 13374–13376 (2002).

29. M. L. Agostini, A. J. Pruijssers, J. D. Chappell, J. Gribble, X. Lu, E. L. Andres, G. R. Bluemling, M. A. Lockwood, T. P. Sheahan, A. C. Sims, M. G. Natchus, M. Saindane, A. A. Kolykhalov, G. R. Painter, R. S. Baric, M. R. Denison, Small-molecule antiviral β-D-N(4)-hydroxycytidine inhibits a proofreading-intact coronavirus with a high genetic barrier to resistance. J Virol 93, (2019).

30. L. F. Cassidy, J. L. Patterson, Mechanism of La Crosse virus inhibition by ribavirin. Antimicrob Agents Chemother 33, 2009–2011 (1989).

31. S. H. Khoo, R. FitzGerald, G. Saunders, C. Middleton, S. Ahmad, C. J. Edwards, D. Hadjiyiannakis, L. Walker, R. Lyon, V. Shaw, P. Mozgunov, J. Periselneris, C. Woods, K. Bullock, C. Hale, H. Reynolds, N. Downs, S. Ewings, A. Buadi, D. Cameron, T. Edwards, E. Knox, I. Donovan-Banfield, W. Greenhalf, J. Chiong, L. Lavelle-Langham, M. Jacobs, J. Northey, W. Painter, W. Holman, D. G. Lalloo, M. Tetlow, J. A. Hiscox, T. Jaki, T. Fletcher, G. Griffiths, Molnupiravir versus placebo in unvaccinated and vaccinated patients with early SARS-CoV-2 infection in the UK (AGILE CST-2): a randomised, placebo-controlled, double-blind, phase 2 trial. Lancet Infect Dis 23, 183–195 (2023).

32. A. B. Evans, C. W. Winkler, K. E. Peterson, Differences in neuropathogenesis of encephalitic California serogroup viruses. Emerg Infect Dis 25, 728–738 (2019).

33. C. A. Schneider, W. S. Rasband, K. W. Eliceiri, NIH Image to ImageJ: 25 years of image analysis. Nat Methods 9, 671–675 (2012).

