## Supplemental Fig. 1 for "*N*^4^-Hydroxycytidine/Molnupiravir Inhibits RNA-Virus Induced Encephalitis by Producing Mutated Viruses with Reduced Fitness"

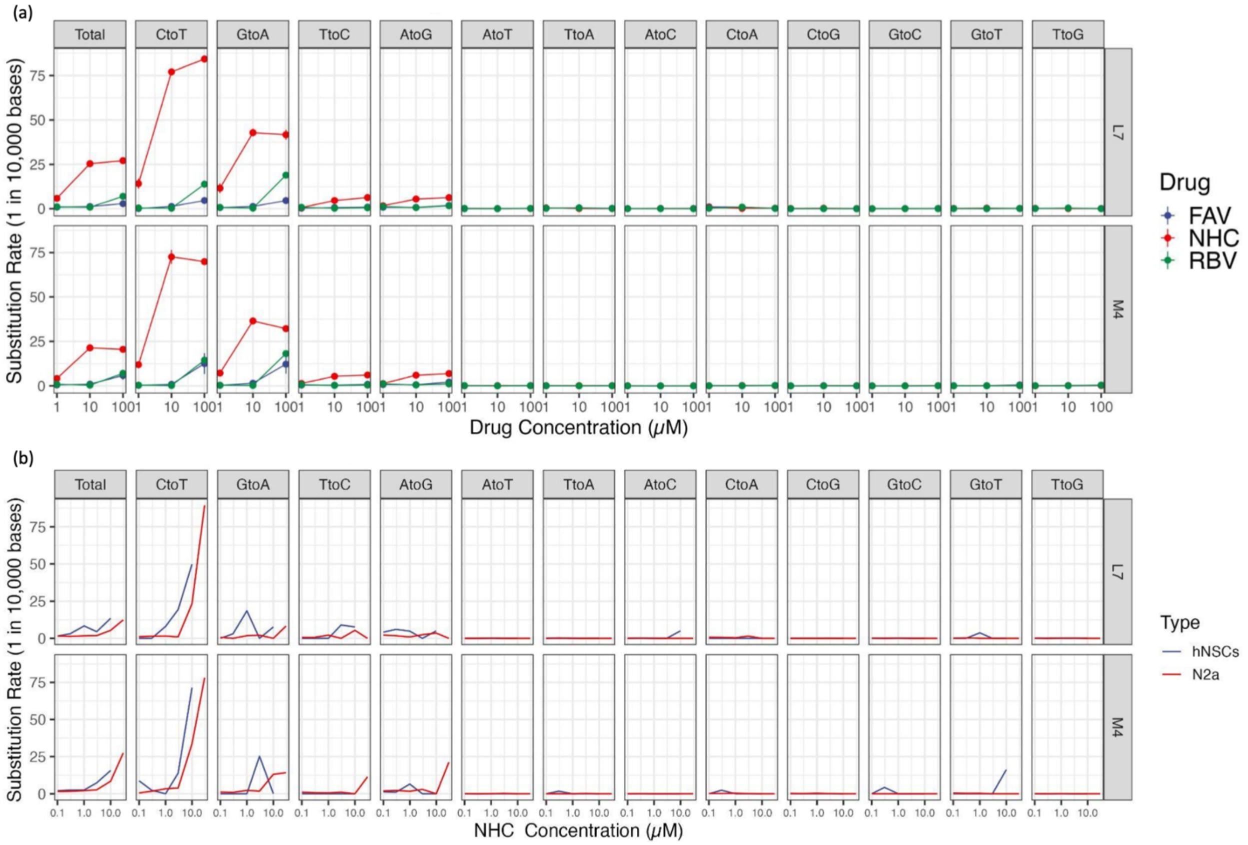


**Supplemental Figure 1**. Mechanism of action of NHC, RBV and FAV in cell models. (a) Substitution rate measured by MPID-NGS in the LACV Vero cell model. (b) Substitution rate measured by MPID-NGS in the LACV Na2 and hNSC cell models.
