## Supplementary material for "*N*^4^-Hydroxycytidine/Molnupiravir Inhibits RNA-Virus Induced Encephalitis by Producing Mutated Viruses with Reduced Fitness": Table S1

**Table S1.** Cytotoxicity of three nucleoside analogs by MTT assay.

| Drugs | CC50 (μM) | |
|---|---|---|
| | Vero cells | hNSCs |
| RBV | 1777.5 | 1264.7 |
| FAV | 3341.2* | 1059.8* |
| NHC | 203.1 | 131.9 |

*$CC_{50}$ of FAV was copied and converted to μM from μg/ml and represent here.
