## Supplementary material for "*N*^4^-Hydroxycytidine/Molnupiravir Inhibits RNA-Virus Induced Encephalitis by Producing Mutated Viruses with Reduced Fitness": Table S2

**Table S2.** LACV MPID-NGS library prep primers.

| | | |
|---|---|---|
| L_PID11 | GTGACTGGAGTTCAGACGTGTGCTCTTCCGATCTNNNNNNNNNNNCAGTTCCTGTGGGTAGAGGATAGG | cDNA primer. Targeting LACV RdRp gene found on the large genomic segment. |
| M_PID11 | GTGACTGGAGTTCAGACGTGTGCTCTTCCGATCTNNNNNNNNNNNCAGTAATCCGATAGATGTCCCAGC | cDNA primer. Targeting LACV medium genomic segment. |
| L_AD | GCCTCCCTCGCGCCATCAGAGATGTGTATAAGAGACAGNNNNAAGGCCAGAAAACGTCAAAG | 1st round PCR forward primer. Targeting LACV RdRp gene found on the large genomic segment. |
| M_AD | GCCTCCCTCGCGCCATCAGAGATGTGTATAAGAGACAGNNNNTCCTGACGTAAAGCTCATCC | 1st round PCR forward primer. Targeting LACV medium genomic segment. |
| ADPT_2a | GTGACTGGAGTTCAGACGTGTGCTC | 1st round PCR reverse primer |
| P1 | AATGATACGGCGACCACCGAGATCTACACGCCTCCCTCGCGCCATCAGAGATGTG | 2nd round PCR forward primer with Illumina adapter sequence |
| Old Nextera | GCCTCCCTCGCGCCATCAGAGATGTGTATAAGAGACAG | Customized sequencing primer |
| Index01 | CAAGCAGAAGACGGCATACGAGAT**CGTGAT**GTGACTGGAGTTCAGACGTGTGCTC | 2nd round PCR reverse indexed primer |
| Index02 | CAAGCAGAAGACGGCATACGAGAT**ACATCG**GTGACTGGAGTTCAGACGTGTGCTC | 2nd round PCR reverse indexed primer |
| Index03 | CAAGCAGAAGACGGCATACGAGAT**GCCTAA**GTGACTGGAGTTCAGACGTGTGCTC | 2nd round PCR reverse indexed primer |
| Index04 | CAAGCAGAAGACGGCATACGAGAT**TGGTCA**GTGACTGGAGTTCAGACGTGTGCTC | 2nd round PCR reverse indexed primer |
| Index05 | CAAGCAGAAGACGGCATACGAGAT**CACTGT**GTGACTGGAGTTCAGACGTGTGCTC | 2nd round PCR reverse indexed primer |
| Index06 | CAAGCAGAAGACGGCATACGAGAT**ATTGGC**GTGACTGGAGTTCAGACGTGTGCTC | 2nd round PCR reverse indexed primer |
| Index07 | CAAGCAGAAGACGGCATACGAGAT**GATCTG**GTGACTGGAGTTCAGACGTGTGCTC | 2nd round PCR reverse indexed primer |
| Index08 | CAAGCAGAAGACGGCATACGAGAT**TCAAGT**GTGACTGGAGTTCAGACGTGTGCTC | 2nd round PCR reverse indexed primer |
| Index09 | CAAGCAGAAGACGGCATACGAGAT**CTGATC**GTGACTGGAGTTCAGACGTGTGCTC | 2nd round PCR reverse indexed primer |
| Index10 | CAAGCAGAAGACGGCATACGAGAT**AAGCTA**GTGACTGGAGTTCAGACGTGTGCTC | 2nd round PCR reverse indexed primer |
| Index11 | CAAGCAGAAGACGGCATACGAGAT**GTAGCC**GTGACTGGAGTTCAGACGTGTGCTC | 2nd round PCR reverse indexed primer |

| | | |
|---|---|---|
| Index12 | CAAGCAGAAGACGGCATACGAGAT**TACA** **AG**GTGACTGGAGTTCAGACGTGTGCTC | 2nd round PCR reverse indexed primer |
| Index13 | CAAGCAGAAGACGGCATACGAGAT**TATG** **GA**GTGACTGGAGTTCAGACGTGTGCTC | 2nd round PCR reverse indexed primer |
| Index14 | CAAGCAGAAGACGGCATACGAGAT**TAGT** **AC**GTGACTGGAGTTCAGACGTGTGCTC | 2nd round PCR reverse indexed primer |
| Index15 | CAAGCAGAAGACGGCATACGAGAT**TACT** **GT**GTGACTGGAGTTCAGACGTGTGCTC | 2nd round PCR reverse indexed primer |
| Index16 | CAAGCAGAAGACGGCATACGAGAT**CATG** **AG**GTGACTGGAGTTCAGACGTGTGCTC | 2nd round PCR reverse indexed primer |
| Index17 | CAAGCAGAAGACGGCATACGAGAT**TATC** **GT**GTGACTGGAGTTCAGACGTGTGCTC | 2nd round PCR reverse indexed primer |
| Index18 | CAAGCAGAAGACGGCATACGAGAT**TCTG** **CA**GTGACTGGAGTTCAGACGTGTGCTC | 2nd round PCR reverse indexed primer |
| Index19 | CAAGCAGAAGACGGCATACGAGAT**TATG** **AA**GTGACTGGAGTTCAGACGTGTGCTC | 2nd round PCR reverse indexed primer |
| Index20 | CAAGCAGAAGACGGCATACGAGAT**ACAG** **CA**GTGACTGGAGTTCAGACGTGTGCTC | 2nd round PCR reverse indexed primer |
| Index21 | CAAGCAGAAGACGGCATACGAGAT**GTGA** **TA**GTGACTGGAGTTCAGACGTGTGCTC | 2nd round PCR reverse indexed primer |
| Index22 | CAAGCAGAAGACGGCATACGAGAT**TATC** **CA**GTGACTGGAGTTCAGACGTGTGCTC | 2nd round PCR reverse indexed primer |
| Index23 | CAAGCAGAAGACGGCATACGAGAT**AATG** **CG**GTGACTGGAGTTCAGACGTGTGCTC | 2nd round PCR reverse indexed primer |
| Index24 | CAAGCAGAAGACGGCATACGAGAT**TAAG** **GC**GTGACTGGAGTTCAGACGTGTGCTC | 2nd round PCR reverse indexed primer |
